# Helpers increase daily survival rate of Southern Lapwing (*Vanellus chilensis*) nests during the incubation stage

**DOI:** 10.1101/179606

**Authors:** Eduardo S. A. Santos, Regina H. Macedo

## Abstract

Cooperative breeding is characterized by reproduction in the presence of helpers. What impact these helpers have on the reproductive success of group members is one of the long-standing questions in the cooperative breeding literature. In cooperative species, helpers are known to provide benefits during multiple stages or at a particular stage of the reproductive cycle. The aim of this study was to investigate whether helpers increased the daily survival rate of nests during the incubation stage in the Southern Lapwing (*Vanellus chilensis*), a crested plover with a cooperative breeding system. Southern Lapwings have a variable mating system, with some breeding groups composed of unassisted pairs, and others that breed in the presence of helpers. Our best supported model indicated a positive effect of the presence of helpers on the daily survival rate of nests, leading to a probability of nest success (i.e., survival until hatching) of 83%, compared to 51% for nests of unassisted pairs. But a null model had a similar model weight as the best supported model and was the second-best model. Our study provides evidence that helpers influence egg survival during the egg-incubation stage, which could influence the fitness of breeders.

## 1 Introduction

A defining characteristic of cooperative breeding systems is the presence of non-breeding helpers (Brown, 1987). The role that these helpers play during the reproduction of group breeders has been the focus of much research over decades (reviewed in Koenig & Dickinson, 2004). How these helpers influence the reproductive success of breeding individuals in their groups, and why they forego direct reproduction while providing help are among the main questions concerning cooperative breeding systems (Emlen, 1991, Dickinson & Hatchwell, 2004). It is commonly thought that helpers are allowed to remain in the group because they positively influence the fitness of the breeding pair. Empirical studies have found evidence of beneficial effects of helping through increased group productivity (e.g., Dias *et al.*, 2015), improved offspring performance (e.g., Brouwer *et al.*, 2012), higher breeder survival (e.g., Russell *et al.*, 2007), or reduction in maternal investment (e.g., Russell *et al.*, 2007, Santos & Macedo, 2011). However, other studies have found little or even contrary evidence of the positive effect of helpers on the productivity of their group’s breeders (e.g., Walters, 1990, Legge, 2000).

The presence of helpers in reproductive groups may cause breeders to adjust their physiology or behaviour. For instance, models predict that breeding females may adjust their clutch size in expectation of the extra parental care that the offspring will receive (differential allocation hypothesis, Burley, 1986, Russell & Lummaa, 2009). On the other hand, theory also predicts the possibility that breeders may reduce their investment into the clutch or eggs, and even reduce the amount of parental behaviour once offspring are born (load-lightening hypothesis, Brown & Brown, 1981, Russell & Lummaa, 2009). In other species, breeders may not be able to plastically adjust their reproductive physiology or behaviour, despite the presence of helpers.

In all of these cases, the presence of helpers is still predicted to increase group productivity and, in addition, helpers may improve the conditions experienced by the offspring, when compared to pairs that breed unassisted. Helpers may, for instance, increase the group’s ability to protect offspring from predators (e.g., Taborsky *et al.*, 2007), or — in birds — improve incubation conditions (e.g., Radford, 2004). Such forms of helping may thus allow initially similar sized clutches/broods — when compared to unassisted breeding pairs — to fare better, leading to greater group productivity. Helping behaviour aimed at the egg incubation stage should yield large benefits, especially to precocial bird species, in which offspring typically feed independently after hatching.

Adaptive explanations for the evolution of helping behavior argue that in some contexts helpers gain higher fitness benefits by helping others than by breeding independently (Koenig & Dickinson, 2016). Helpers gain indirect fitness benefits when they provide helping behavior that is of relatively high value and increase the fitness of individuals to whom they are genetically related. The value of parental care in birds is thought to be relatively higher in altricial than in precocial species, as in the latter case the offspring are often very independent after hatching. Given that helpers gain more indirect benefits when the value of parental care is higher, one should expect helping behavior to be more common in altricial than in precocial bird species. Yet, cooperative breeding occurs in precocial species that exhibit costly parental care (see Walters, 1982), which leads to the question of how much benefit helpers provide in such cases.

The aim of this study was to investigate whether helpers increase the daily survival rate of nests during the incubation period in the cooperative breeding system of the Southern Lapwing, *Vanellus chilensis*. Southern lapwings are precocial plovers that occur throughout South America. During the breeding season, pairs may breed unassisted or in the presence of helpers-at-the-nest (Walters & Walters, 1980, Saracura *et al.*, 2008, Santos & Macedo, 2011). The fact that cooperative breeding in plovers is rare makes the Southern Lapwing a unique study system in which to address questions of cooperative behavior.

## 2 Materials & Methods

### 2.1 Study area, species and general methods

We conducted fieldwork during one breeding season (Aug–Dec 2007) in two neighbourhoods of the city of Brasília, Brazil (15°81’S 47°87’W and 15°91’S 47°94’W). Both areas present extensive lawns that are used by Southern Lapwings as breeding territories (Santos & Macedo, 2011). We searched for nests visually within the territories of pairs and cooperative groups. Once a nest was located, we monitored it every 2–4 days, until the chicks hatched or the nest was depredated.

During each visit, we recorded the number and condition of eggs within each nest. The fate of a nest was considered successful if ≥ 1 egg hatched (i.e., if recently-hatched chicks were observed within the monitored nest scrape, coupled with a sequential decrease in the number of eggs at a nest). A nest was considered to have failed if it was abandoned (eggs were cold), depredated (signs of predation such as broken eggshells), or failed to hatch (hatching never occurred even though parents continued to incubate).

To determine whether lapwing pairs bred with or without helpers, we monitored the territories to determine the maximum number of adult individuals that engaged in parental activities during at least two consecutive nest visits. we considered parental activities to be either incubation of the eggs or nest defence against predators (or researchers). We did not consider conspecific neighbors and floaters that joined the focal group during mobbing instances as part of the breeding groups. Only individuals that were inside the focal group’s territory at the beginning of nest defense behaviors were considered when determining group size. None of the monitored breeding groups changed their size over a period of two breeding seasons (2007–2008; ESAS personal observation). Six (6) cooperative groups were composed by a trio of adults, while the other two groups were composed of five (5) adults.

### 2.2 Analyses

We estimated daily nest survival with the nest survival model in *MARK* (White & Burnham, 1999, Dinsmore *et al.*, 2002) using the *RMark* package (Laake, 2013) as an interface in *R* 3.3.0 (R Core Team, 2016). We were interested in the effect of the breeding group composition on DSR, thus we used the type of breeding group composition (pair or cooperative group) as a categorical predictor in the DSR model. We treated breeding group composition as a categorical predictor with two levels, because group size in the cooperative groups showed little variation. We also fit models with: (1) a continuous time trend, (2) nest age, (3) nest age and group type, and (4) a constant DSR model. We then used Akaike’s Information Criterion (AIC) (Burnham & Anderson, 2002), adjusted for small samples (AICc), to rank all the models. We calculated DSR based on regression coefficients from the most-supported model (i.e., the one with the lowest AICc value).

Based on the best model, we calculated the total nest survival using a 28-day incubation period (Saracura, 2003), and calculated the 85% confidence interval (CI). We used an 85% CI to interpret the effect of breeding group type of total nest survival because it allowed for more congruence (Arnold, 2010).

## 3 Results

We monitored a total of 25 and 11 nests in 2007 from monogamous pairs and cooperative groups, respectively. Five nests were removed from analyses because they were abandoned or depredated and we did not have the dates in which the events took place, thus preventing us from using these data in the analyses. Of the remaining nests (23 from pairs and 8 from cooperative groups), 22 were successful and 9 were unsuccessful.

The top model included type of breeding group as a predictor, with a model weight of 0.245 (Table 1). However, the null model had a similar model weight of 0.240 and was the second-best model (Table 1). The nest survival estimates of the best model showed that nests cared for by pairs had a tendency to have lower survival than nests of groups, but the confidence interval of the slope did overlap zero (*β*_pairs_ = −1.288 [85% CI: −2.812 to 0.235]). The estimated daily survival rate of nests tended by cooperative groups was higher than that of nests tended by monogamous pairs (DSR_pairs_ = 0.976 [85% CI: 0.962 to 0.985]; DSR_cooperative groups_ = 0.993 [85% CI: 0.972 to 0.998]). The probability of success (DSR raised to an exponent of 28 (incubation days)) of a nest tended by cooperative groups is 0.832 (85% CI: 0.276 to 0.974), while the probability of success of nests tended by pairs is 0.513 (85% CI: 0.281 to 0.709).

**Table 1:** Model selection based on Akaike’s Information Criterion adjusted for small sample size (AICc) of daily survival rates (DSR) of Southern Lapwing nests during the incubation stage.

| Model | Parameters | AICc | $\Delta AICc$ | Weight | Deviance |
| --- | --- | --- | --- | --- | --- |
| $S(\sim \text{GroupType})$ | 2 | 78.48 | 0 | 0.25 | 74.46 |
| $S(\sim 1)$ | 1 | 78.52 | 0.04 | 0.24 | 76.52 |
| $S(\sim \text{NestAge} + \text{GroupType})$ | 3 | 80.48 | 2.0 | 0.09 | 74.43 |
| $S(\sim \text{NestAge})$ | 2 | 80.52 | 2.03 | 0.09 | 76.49 |
| $S(\sim \text{Time})$ | 2 | 80.54 | 2.05 | 0.09 | 76.51 |

## 4 Discussion

The aim of this study was to investigate if helpers increased the daily survival rate of nests in the Southern Lapwing. We confirmed that the daily survival rate of nests tended by cooperative groups was higher than that of unassisted pairs, but note that the 85% CIs overlapped. Yet, when considering a 28-day nesting period, our findings suggest that the presence of helpers in breeding groups increases the probability of nest success compared with nests of unassisted pairs. Our estimates yield an 83% probability of nest success for those nests tended by cooperative groups, while a probability of 51% for nests of unassisted pairs.

Several studies have investigated whether the presence of helpers in cooperative breeding groups leads to increased reproductive success of breeding group members (reviewed in Koenig & Dickinson, 2004, 2016). One of the ways in which helpers may be able to generate positive effects on the reproductive success of group members is through assistance during the egg incubation stage. For instance, helpers may act as sentinels and alert incubating individuals of the presence of potential nest predators (e.g., Alves, 1990). Additionally, helpers may take over the incubation of the eggs, which, again, may lead to more stable egg-development conditions (e.g., Dias *et al.*, 2013).

Interestingly, several empirical studies of cooperative breeding birds have failed to find a biologically meaningful effect of helpers on the probability of nest survival (e.g., Magrath & Yezerinac, 1997, Blackmore & Heinsohn, 2007, Manica & Marini, 2012). One difference between the studied species and the Southern Lapwing is that they are altricial, while Southern Lapwings have precocial development. A potential explanation for this difference, and which has not been empirically tested, is that precocial species typically tend to have longer incubation periods than altricial species. For instance, the average incubation period of the altricial White-banded Tanager, *Neothraupis fasciata*, lasts 13 days (Manica & Marini, 2012), while the incubation period of precocial Southern Lapwings lasts 28 days. Thus, the longer incubation period may be more exposed to predators or environmental variability, which in turn could make additional investment of helpers more beneficial, or at least, more detectable. Another, explanation for the difference in the benefits of helpers between precocial and altricial species is that care in precocial species could be more concentrated during the egg-stage, while in altricial species very little help could be directed at the egg-stage.

Among precocial species, Walters (1982) argued that Southern Lapwings exhibited the highest time costs of parental care among some of the studied species and suggested that costly parental care was linked to the evolution of cooperative breeding. We have shown here that helpers increased the probability of nest success, which could lead to higher fitness benefits in cooperative groups if more offspring, from these groups, fledge and are recruited (not investigated in this study). Thus, it is possible that helpers in the Southern Lapwing gain sufficient indirect fitness benefits to compensate independent reproduction. A comprehensive investigation of fitness benefits and the value of parental care in Southern Lapwings would be a valuable addition to our understanding of the evolution of cooperative breeding in precocial species.

Overall, our data suggest that helpers in Southern Lapwings increase the probability of success of nests. The probability that a nest will be successful is an especially important component of the reproductive success in precocial species, in which offspring are considerably independent after hatching.

## 5 Acknowledgments

We would like to thank Pedro Diniz and Eduardo Carvalho for their valuable help during data collection. We also thank two anonymous reviewers for comments and suggestions on an earlier version of this manuscript.

## 8 Author contributions

Eduardo S. A. Santos conceived and designed the study, collected and analysed the data, wrote, and reviewed drafts of the manuscript.

Regina H. Macedo reviewed and edited the original manuscript.

## 9 Field study permission

This study was conducted with the proper approval from the Instituto Brasileiro do Meio Ambiente e dos Recursos Naturais Renováveis (IBAMA).

## 10 Data accessibility

Data for this article can be found online at the figshare repository.

